# Diversification, loss, and virulence gains of the major effector AvrStb6 during continental spread of the wheat pathogen *Zymoseptoria tritici*

**DOI:** 10.1101/2024.10.12.618020

**Authors:** Ana Margarida Sampaio, Sabina Moser Tralamazza, Faharidine Mohamadi, Yannick De Oliveira, Jérôme Enjalbert, Cyrille Saintenac, Daniel Croll

**Affiliations:** Laboratory of Evolutionary Genetics, Institute of Biology, University of Neuchâtel, 2000 Neuchâtel, Switzerland; Arvalis - Institut du Végétal, Station expérimentale, 91720 Boigneville, France; Université Paris-Saclay, INRAE, CNRS, AgroParisTech, GQE-Le Moulon, 91190, Gif-sur-Yvette, France; Université Clermont Auvergne, INRAE, GDEC, 63000, Clermont-Ferrand, France

## Abstract

Interactions between plant pathogens and their hosts are highly dynamic and mainly driven by pathogen effectors and plant receptors. Host-pathogen co-evolution can cause rapid diversification or loss of pathogen genes encoding host-exposed proteins. The molecular mechanisms that underpin such sequence dynamics remains poorly investigated at the scale of entire pathogen species. Here, we focus on *AvrStb6*, a major effector of the global wheat pathogen *Zymoseptoria tritici*, evolving in response to the cognate receptor *Stb6*, a resistance widely deployed in wheat. We comprehensively captured effector gene evolution by analyzing a global thousand-genome panel using reference-free sequence analyses. We found that AvrStb6 has diversified into 59 protein isoforms with a strong association to the pathogen spreading to new continents. Across Europe, we found the strongest differentiation of the effector consistent with high rates of *Stb6* deployment. The *AvrStb6* locus showed also a remarkable diversification in transposable element content with specific expansion patterns across the globe. We detected the *AvrStb6* gene losses and evidence for transposable element-mediated disruptions. We used genome-wide association mapping data to predict virulence emergence and found marked increases in Europe, followed by spread to subsequently colonized continents. Finally, we genotyped French bread wheat cultivars for *Stb6* and monitored resistant cultivar deployment concomitant with *AvrStb6* evolution. Taken together, our data provides a comprehensive view of how a rapidly diversifying effector locus can undergo large-scale sequence changes concomitant with gains in virulence on resistant cultivars. The analyses highlight also the need for large-scale pathogen sequencing panels to assess the durability of resistance genes and improve the sustainability of deployment strategies.

**Author summary:** Interactions between plants and their specialized pathogens are often mediated by a sophisticated molecular dialogue. Effectors produced by pathogens serve to manipulate the host but may also be used by the host to trigger defence mechanisms upon recognition. Deploying plants carrying a resistance gene against a specific effector could lead to rapid adaptation in the pathogen. Here, we unraveled such dynamics at the scale of the global distribution range of the fungal wheat pathogen *Zymoseptoria tritici*. The effector is encoded by the gene *AvrStb6* located in a polymorphic region of a chromosome near the telomere. We find selfish elements (*i.e.* transposable elements) repeatedly inserted nearby the gene, which has likely facilitated the rapid sequence evolution. The effector diversified among continents, and we could predict that the sequence changes likely helped escape recognition by the host receptor. Our study provides one of the most comprehensive views how a crop pathogen diversified a major effector in response to host resistance factors. Such studies facilitate devising more durable deployment strategies of host resistance in order to maintain crop yield.

## Introduction

Interactions between plant pathogens and their host is a highly dynamic process, mainly mediated by pathogen effectors (*i.e.* avirulence factors, *Avr*) and resistance (*R*) plant genes. Effector genes encode secreted molecules capable of modulating host plant metabolism or suppress plant immune responses, thereby are crucial for successful host infection (1). In turn, *R* genes encode proteins that can recognize specific pathogen effectors, particularly those encoded by *Avr* genes, subsequently triggering an immune response (2,3). In this gene-for-gene model, the presence of *R* genes imposes selection pressure on pathogens carrying recognized effectors (Stukenbrock and McDonald, 2009). This favors effector mutations preventing recognition by purging avirulent protein variants (4,5). Mechanisms include transposon insertions (6), repeat-induced mutations (RIP) (7) or even complete loss of recognized effector. Being beneficial, those mutations often spread rapidly through the pathogen population (8).

The dynamics of effector evolution are often influenced by their chromosomal sequence environment. They can be located in either accessory chromosomes enriched in repetitive sequences (9) or transposable element (TE) rich core chromosome compartments (9,10). Repeat proximity facilitates effector sequence diversification and, hence, increases mutations available for adaptation to an evolving host (11,12). Such rapid virulence evolution was described in the rice pathogen *Magnaporthe oryza* due to effector localization in highly repetitive subtelomeric region. Mutations in *AVR-Pita* effectors, such as point mutations, insertions and deletions, have enabled the fungus to evolve avoiding triggering immune response (13). Furthermore, this effector has undergone multiple translocations associated with virulence evolution (14). The transposition of TEs can disrupt effector coding sequences or alter their regulation. In *M. oryzae*, the insertion of a Mg-SINE TE in the *AvrPi9* gene led to loss-of-function of the effector (15), while in *Verticilium dahlia*, a TE insertion inactivated the *Ave1* effector (16), both probably enabling the pathogen to escape host recognition. In addition, repeat-induced point (RIP) mutations, a defense mechanism against TEs, can impact effector diversification and enhance virulence. In the fungal pathogen *L. maculans*, the emergence of virulence alleles in *AvrLm1* was driven by RIP mutations, resulting in a non-functional locus and host resistance breakdown (17). Furthermore, leakage of RIP into neighboring regions contributed to the diversification of effector genes while retaining functionality (7). Taken together, effector gene diversification and TE dynamics of the surrounding regions are key factors to assess the evolutionary potential of the pathogen. However, how the effectors evolved in response to strong host selection pressure across and within species remains poorly understood.

*Zymoseptoria tritici*, the causal agent of septoria tritici blotch (STB), is one of the major fungal foliar diseases of wheat-growing areas worldwide (18). The spread of the pathogen has been tightly associated with the origins of wheat cultivation (19). With its center of origin located in the Middle East, *Z. tritici* initially colonized North Africa and Europe, and later migrated to the Americas and Oceania (20). Population genomic analyses identified eleven well-supported genetic clusters with most being restricted to individual continents. In the Middle East, two distinct clusters distinguished isolates from Iran and Israel. Isolates collected in Northern Africa and Europe were also represented by two different clusters, while Australian and New Zealand isolates were grouped into three clusters. North American *Z. tritici* populations grouped in two clusters along a North-South separation, while in South America two clusters split pathogen diversity along the Andes (20). Even though continental divisions reflect historic restrictions in gene flow, a significant fraction of isolates mismatched the prevalent genetic cluster on the continent consistent with recent migration events. Europe was the strongest source for such events across continents (20). Beyond changes in genetic diversity, *Z. tritici* has also undergone shifts in transposable element (TE) content, with the most recently colonized areas (Americas and Oceania) showing higher numbers of genome-wide TE insertions and incipient expansions in genome size likely associated by the loss of RIP activity (20,21). TEs cover 16.5-24% of the genome and are often located near genes involved in host-interactions (22). TEs activity can also be directly associated with effector gene expression and virulence (23–25). The best understood effector gene influenced by TE activity is *Avr3D1* with the insertion and subsequent silencing of a TE contributing to downregulation of the recognized effector and evasion of host recognition (24). Additionally, specific amino acid changes in this effector sequence have also been shown to lead to evasion of recognition (23).

The first effector to be cloned in *Z. tritici* was *AvrStb6* (26,27), which plays a dominant role in immunity evasion due to the prevalence of the corresponding resistance gene. The effector is encoded in a highly polymorphic subtelomeric region of chromosome 5 surrounded by TEs (8,27,28). *AvrStb6* is recognized by the plant wall-associated receptor-like kinase *Stb6* (29), present in many wheat cultivars worldwide, as it has been frequently used in breeding programs to control septoria tritici blotch (STB), though its usage varies across different regions (30,31). Given the broad deployment of *Stb6*, *Z. tritici* is expected to experience significant host selection pressure to escape recognition. Indeed, *AvrStb6* shows high haplotype diversity across the world (32–34) with a likely absence of the originally described avirulent isoform among recently collected isolates, consistent with efficient counter-selection against avirulent haplotypes. However, no loss of *AvrStb6* has been documented yet, which was interpreted as evidence for essential but not yet known role of *AvrStb6* (Brunner & McDonald, 2018). Given the high plasticity of the subtelomeric region surrounding *AvrStb6* (8), active TEs could continue to reshape sequence diversity in extant populations.

To address how *AvrStb6* diversification occurred during continental spread, we aimed to provide a large-scale population-genomics informed view how *AvrStb6* and the surrounding regions evolved in response to varying selection pressure imposed by past release of *Stb6* in wheat varieties. For that, we recapitulated *AvrStb6* evolution in a thousand-genome panel of *Z. tritici* covering key regions associated with the historical dissemination of wheat cultivation. We combined reference-genome data with short-read sequencing to validate key insights about the evolution of the locus and tracked insertion dynamics of TEs using a newly established high-quality TE library. To connect *AvrStb6* evolution to the deployment of cognate wheat cultivars, we first used genomic prediction to assess virulence trait evolution across the global dataset and, second, tracked the gain of virulence across a European country in conjunction with monitoring wheat cultivar deployment.

## Methods

### *Z. tritici* isolates collection

We performed *AvrStb6* locus analyses on a global collection of *Z. tritici* comprising 1035 strains analyzed by Feurtey et al. (20) (Figure 1A; Supplementary Table S1). *De novo* draft assemblies were produced using the software SPAdes v3.14.1 (35) using trimmed and filtered reads from Trimmomatic v0.39 (36). To ensure acceptable *de novo* assembly qualities, we used QUAST to calculate assembly metrics (37). We analyzed single nucleotide variants (SNVs) called by Feurtey et al. (20) to estimate genetic subdivision across the worldwide distribution range revealing 11 genetic clusters (Middle East – Israel, Middle East – Iran, Middle East – North-Africa, North America – USA, North America – North, Oceania – Australia, Oceania – Tasmania, Oceania – New-Zealand, Europe, South America – East, and South America – West), closely tracking the historic expansion of wheat cultivation around the world (20). The genome of the closest sister species (*Z. pseudotritici*) (38) was included for phylogenetic analyses of *AvrStb6*. Additionally, we included reference-quality genome assemblies of 19 strains representative of the global genetic diversity of the species (22) for *AvrStb6* loss-of-function analyses.

**Figure 1:**
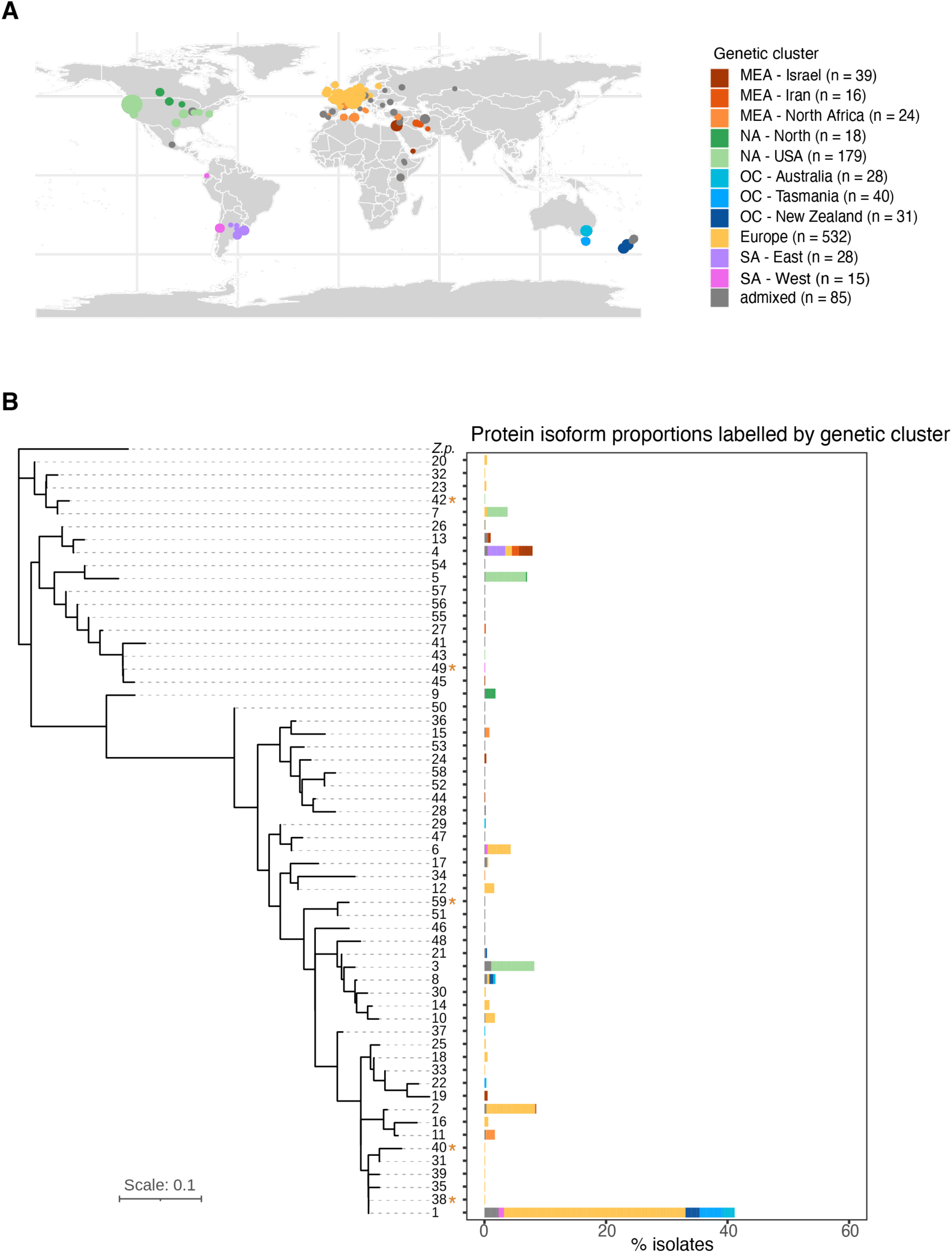
Global panel of *Zymoseptoria tritici* isolates sampled across continents and AvrStb6 diversification. A) Geographical distribution of the 1035 analyzed isolates colored by the 11 assigned genetic clusters (20). Circle sizes are proportional to the population size sampled per location. B) Maximum likelihood phylogenetic tree of AvrStb6 protein isoforms rooted by the *Z. pseudotritici* (*Zp*) closest sister species isoform. Truncated isoforms with premature stop codon are marked with an asterisk. The barplot shows the frequency of isoforms and their composition based on genetic clusters assigned to the isolates.

### *AvrStb6* homologs and phylogenetic analyses

We used draft genome assemblies produced by SPAdes to search for homologs of *AvrStb6* among all *Z. tritici* and *Z. pseudotritici* isolates using the *AvrStb6* IPO323 sequence as query for BLASTn analyses (39). Blastn hits were filtered for a minimum identity of 90%, minimum length (bp) match of 90% (> 330 bp), and maximum *e*-value of 10^-5^. *AvrStb6* haplotype sequences were extracted from draft assemblies based on BLASTn hit coordinates. Subsequently, the coding region sequences were aligned using MAFFT v7.520 (40) and translated to amino acid sequences using EMBOSS Transeq (41). Translated coding sequences were re-aligned using MAFFT v7.520 and a maximum likelihood tree built with 1000 bootstrap replicates (MEGA11, (42)). The tree was rooted using the Z. *pseudotritici* AvrStb6 sequence.

### Transposable elements in *AvrStb6* region

To detect TE insertions near *AvrStb6*, all Illumina scaffolds having *AvrStb6* homologs were annotated with the curated *Z. tritici* TE consensus sequences (43) using RepeatMasker v4.0.9_p2. Annotated elements shorter than 50 bp were filtered out. Since the number of TEs can vary according to the scaffold length, only the closest TEs up and downstream of *AvrStb6* have been considered. TE superfamilies present in more than 50 isolates were considered as frequent.

### Assessing *AvrStb6* loss-of-function variants

To explore possible *AvrStb6* deletions, we first analyzed the 19-reference genome panel (22). Specifically, for the isolate Arg00 (one of the genomes included in the panel), we analyzed raw PacBio read mapping to the *Z. tritici* IPO323 reference genome (44) using Minimap2 v2.17 (45). Mapped reads were assembled using Canu v2.2 (46). All assemblies were performed with an estimated genome size of 40.7 Mb (--genomeSize), error rates of 0.045 (--correctedErrorRate), minimal read length of 500 (--minReadLength) and maximum threads of 32 (--maxThreads). Alignments between the new Arg00 assemblies and IPO323 reference genome in the *AvrStb6* surrounding region were visualized using IGV v2.16.1 software (47). Synteny was plotted with genoplotR v0.8.11 (48) for the 19-reference genome panel, using *Z. tritici* improved gene and TE annotation (43,49).

Possible *AvrStb6* deletions in the thousand-genome panel have been explored by inspecting copy number variation (CNV) calling and read count analyses of the *AvrStb6* genomic region. CNV calling was performed in the global population *de novo* draft assemblies (n = 1109) from Feurtey et al. (50). GATK CNV caller v4.1.9.0 (51) with recommended parameters on aligned BAM files has been used, with CNV interval set to 1-kb window with no overlap. Filtering criteria included GC content (min = 0.1 and max = 0.9) and removal of extremely low and high read counts (50).

### Genomic prediction for *AvrStb6*-mediated virulence

To predict *AvrStb6*-mediated virulence expressed by isolates included in the thousand-genome *Z. tritici* panel, we performed genomic prediction analysis based on the genomic best linear unbiased prediction (gBLUP) method implemented in GAPIT v3.4.0 (52). For this, we retrieved matching phenotype-genotype datasets established for genome-wide association mapping (GWAS). We focused phenotypic data on a French collection of 103 *Z. tritici* isolates, which have been previously used to map *AvrStb6* by GWAS (27). Phenotypes originally collected for this dataset included the percentage of green leaf area (G), percentage of necrotic leaf area (N), and percentage of leaf area containing pycnidia spores within the inoculated area (S). Three wheat cultivars carrying the resistance gene *Stb6* (“Cadenza”, “Shafir”, and “Caphorn”) were used for independent phenotyping assays. As genotype input, we tested two types of SNP sets: a SNP matrix covering the *AvrStb6* region and 1000 bp up- and downstream of the effector region, and a SNP matrix covering the entire genome. The SNP matrices were generated by filtering for biallelic SNPs (option “M2”) and a minor allele frequency of 5% (-q 0.05: minor) using BCFtools v1.5 (53). The kinship matrix was calculated using plink v1.90 (54) based on genome-wide SNPs. We used vcftools (55) “--thin 1000” to randomly retain only 1 SNP for every 1 kb interval as an input dataset for the calculation of the kinship matrix.

Given that input leaf area phenotypic trait values were measured in percentage (ranging from 0-100%), any phenotypic prediction values falling below 0 and above 100 were adjusted to 0 and 100, respectively. Pearson correlation coefficients between phenotypic traits measured in the French collection versus predicted phenotypic values in the same population were calculated in R v4.3.2 using the *cor* function. Correlation coefficients were used to select the best combination of cultivars and traits from the 9 possible pairings, as well as SNP matrix for performing genomic predictions.

### *Stb6* wheat deployment in France and *Stb6* genotyping

Wheat cultivar deployment data for France were retrieved from the DiverCILand database (https://wheat-diverciland.moulon.inrae.fr/). DiverCILand aims to monitor varietal diversity at various scales and gathers data on the deployment areas of registered bread wheat varieties for 17 European countries. Depending on the country, the data covers 5-30 years of deployment history either at the county or national scale. The data on cultivar deployment in France were produced by FranceAgriMer and covers the period 1981-2018.

*Stb6* genotyping of French cultivars was performed using KASP genotyping assays following manufacturer instructions (LGC Genomics®) and diagnostic markers cfn80047 and cfn80050 (56). The run cycle and data analysis were performed on the LightCycler^®^ 480 Real-Time PCR System (Roche Life Science). Knowledge on *Stb6* allelic versions were also retrieved from the genotyping data of 4506 wheat accessions (57) using diagnostic marker AX-89415184 (58). The percentage of cultivars carrying *Stb6* resistance or susceptible alleles per year was calculated on 1327 French cultivars having both cultivar deployment and genotyping information.

Predicted phenotypic values of 211 French *Z. tritici* strains, collected between 1999 and 2016 were inspected in detail. To determine whether the phenotypic traits observed in the French collection could be significantly explained by the frequency of Stb6-resistant allele deployment in wheat cultivars across France, a linear regression analysis was performed using the *lm* function in R v4.3.2.

## Results

### *Global AvrStb6* genetic diversity

The spread of the fungal wheat pathogen *Z. tritici* from its origin in the Fertile Crescent to other continents has been assessed based on a global panel of >1000 sequenced genomes (20). Single nucleotide variant (SNVs) analyses revealed clear genetic subdivisions across the global distribution range with 11 genetic clusters closely tracking the historic spread of wheat cultivation across the world. Multiple clusters were detected in populations near the centre of origin of wheat cultivation (Middle East – Israel and Iran), regions with more recent wheat cultivation areas showing distinct genetic clusters including Europe and North Africa. The most recent introductions occurred over the last centuries (North and South America, and Oceania). Wheat cultivars carrying the *Stb6* resistance genes are globally distributed (31). To unravel the evolutionary trajectory of *AvrStb6* in *Z. tritici*, we analyzed the thousand-genome panel for *AvrStb6* gene variants (Figure 1A).

We searched draft genome assemblies generated for 1035 isolates sampled across the world (20) for homologs of *AvrStb6*. We identified 1001 assemblies containing single matching homologs, excluding sequences of potentially truncated *AvrStb6* copies. The draft assemblies presented reasonable contiguity with the N50 (length of the shortest contig for which longer length contigs cover at least 50% of the assembly) ranging between 2215 and 215,440 bp (Supplementary Table S2). Intact *AvrStb6* copies were recovered ranging from 361-365 bp in length representing a total of 103 nucleotide sequence haplotypes. The encoded protein sequences ranged from 81-82 amino acids for 59 distinct isoforms (Supplementary Table S3). The most frequent isoform, labelled here as isoform 1 matches isoform I02 identified in previous work (33) (Supplementary Table S3). Isoform 1 was shared by 41% of the collected strains. On the contrary, 49 isoforms were each represented by less than 1% of the analyzed isolates. We found no correlation between isoform frequencies and genetic cluster identity (Figure 1B). Three isolates from the European cluster, and two isolates from the American clusters (North America – USA and South America – West) carried a premature stop codon (Supplementary Table S4). The stop codon position was variable and ranged from the 5^th^ to the 46^th^ amino acid position. Nevertheless, amino acid sequences after the premature stop codon remained mostly conserved compared to the reference genome IPO323 *AvrStb6* haplotype (isoform 6).

We reconstructed the phylogenetic relationships among isoforms to recapitulate the effector diversification. Isolates from Middle East (Israel and Iran) carried isoforms similar to the AvrStb6 isoform recovered from the closest sister species *Z. pseudotritici*. On the contrary, isoform 1, the most frequent isoform in European and Oceanian clusters, was the most divergent isoform to *Z. pseudotritici*, showing that a major transition in *AvrStb6* haplotypes occurred following European colonization (Figure 1B). Additional European and Oceanian isoforms, along with Northern African isoforms, clustered closely together with the most divergent isoform. Isolates from the North America – USA cluster presented the most distinct set of isoforms, with two isoforms clustering together with Middle East isoforms, while another isoform revealed to be more divergent. South America – East isolates presented the same isoform as most of the Middle East isolates (isoform 4), while South America - West isolates revealed to be more divergent (Figure 1B). The five protein isoforms with premature stop codons were found across all major branches of AvrStb6 diversification (Figure 1B). Altogether, these findings indicate that *AvrStb6* diversification occurred most prominently in Europe and by this likely impacting effector trajectories in subsequently colonized continents.

### Transposable element dynamics at the *AvrStb6* locus

TEs tend to be located closer to effector genes compared to other genes in the genome across diverse fungal pathogens. Hence TEs, have a significant potential to act as regulators of effector genes. *AvrStb6* is located in a gene-poor and TE-rich region close to a telomeric end of chromosome 5. To investigate patterns of TE dynamics near *AvrStb6,* we analyzed contigs encoding *AvrStb6* among different isolates to screen for the presence of TE sequences. Upstream of *AvrStb6*, the two most frequently inserted TE superfamilies included a miniature inverted-repeat transposable element (MITE; ZymTri_2023_family_1310) and an unclassified low-copy TE (ZymTri_2023_family_1288), with the latter one found in 796 (79.5%) of isolates. In contrast, populations sampled near the centre of origin (Middle East, Israel and Iran) as well as the South American cluster (SA-East) carried most frequently a MITE (Figure 2A). Isolates from both the European and Oceanian clusters (Australia, Tasmania and New Zealand) carried the unclassified TE most frequently (Figure 2A). Remarkably, all Oceanian clusters carried exclusively the unclassified TE near *AvrStb6*. The MITE ZymTri_2023_family_1310 was at 163 bp from the start of the coding sequence in all the 170 isolates among different population clusters where this TE has been found (Figure 2B and C) consistent with a single, recent insertion event. This TE is also the most detected TE close to *AvrStb6* (Figure 2B and C; Supplementary Table S5). Downstream of *AvrStb6*, we identified distinct insertions by TEs from different superfamilies: LINE retrotransposons (ZymTri_2023_family_1222, 148, and 605), retrotransposons LTR/Gypsy (ZymTri_2023_family_1335, 243, 607, and 981), as well as unclassified TEs (ZymTri_2023_family_1299, 1473, 250, 363, 697, 795). Downstream of *AvrStb6*, inserted TEs were at variable distance to *AvrStb6* (Figure 2B). A single isolate (STnnJGI_SRR7073594, from North America - North) carried an unclassified TE (ZymTri_2023_family_363) just at 8 bp downstream of *AvrStb6*, while 21 isolates from the North America - USA cluster carried as the closest downstream TE a retrotransposon LTR/Gypsy 7339 bp away from *AvrStb6* (Figure 2B; Supplementary Table S5).

**Figure 2:**
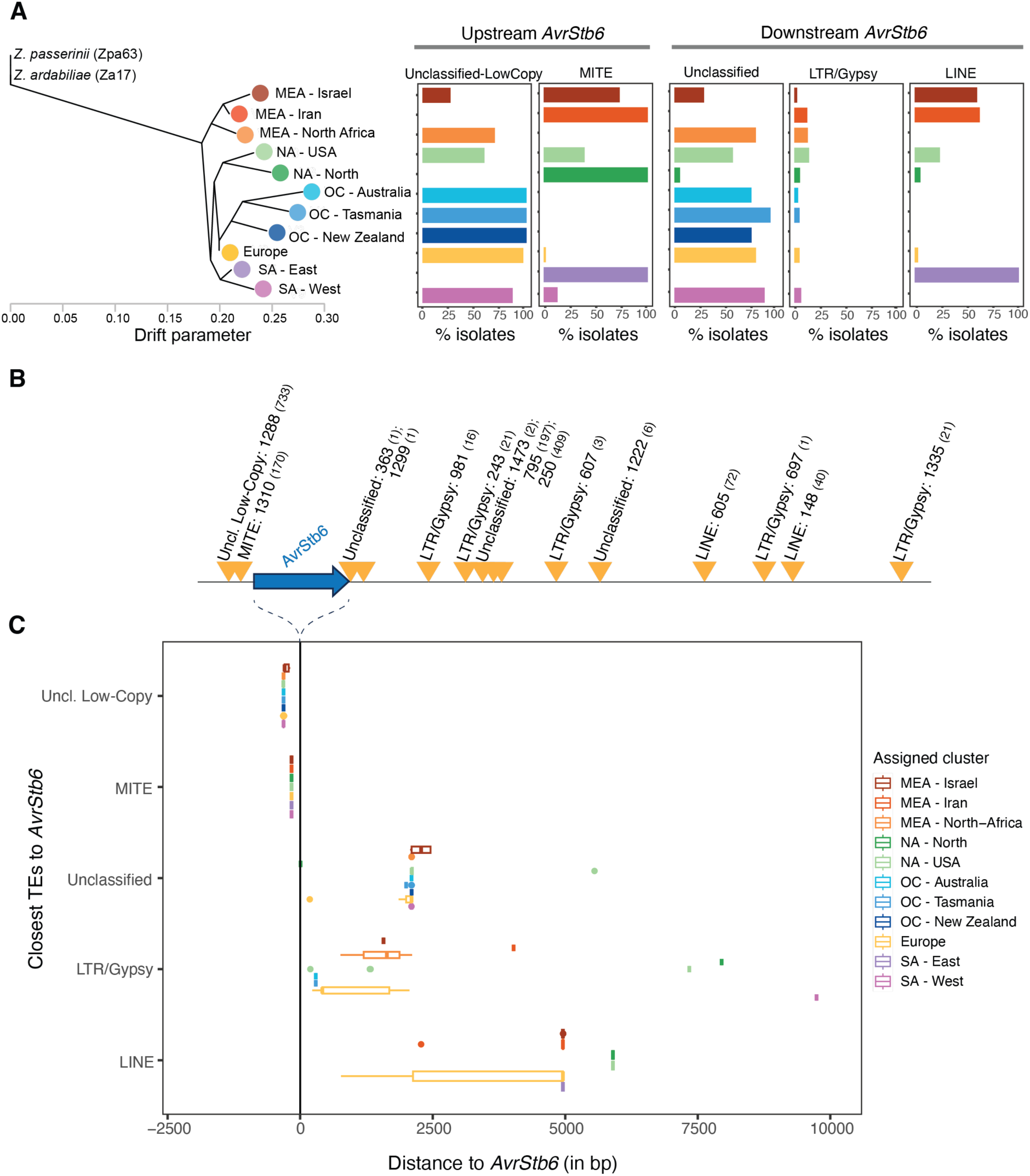
Insertion frequencies of transposable elements (TEs) near *AvrStb6* in the thousand-genome panel. A) TE insertion frequencies for the most frequent TE superfamilies according to the genetic clusters assigned to the isolates. The population tree was generated based on Feurtey et al. (20). B) Schematic representation illustrating the average distance of TE copies grouped by superfamilies. The number after TE superfamily identifiers represents the family number and the number within parentheses represents the number of *Z. tritici* isolates having the respective TE. C) Variation in distance between specific TEs and *AvrStb6* among isolates. Distance variation is summarized by TE superfamily (or unclassified TEs) for each genetic cluster. The line at zero basepairs represents the *AvrStb6* position. Unclassified low-copy TE refers to ZymTri_2023_family_1288; MITE refers to ZymTri_2023_family_1310; LTR/Gypsy refers to ZymTri_2023_family_1335, 243, 607, and 981; Unclassified TE refers to ZymTri_2023_family_1299, 1473, 250, 363, 697, 795; and LINE refers to ZymTri_2023_family_1222, 148, and 605.

Overall, TEs downstream were found at larger and more variable distances to *AvrStb6* compared to upstream TEs (Figure 2B and C; Supplementary Table S5). Furthermore, both close up- and downstream TEs were partially degraded suggesting that TEs were inactivated likely in recent history. Populations from the Middle East (Israel and Iran) both carry LINE retrotransposons as the most frequent TE (Figure 2A). Regions outside of the centre of origin largely carried unclassified TEs except for the South America (East) cluster, which is sharing TE patterns with isolates from the centre of origin. In conjunction, the TE insertion analyses show that TE associations with *AvrStb6* underwent significant shifts as the pathogen spread from the Middle East to North-Africa, Europe and later introductions to the Americas and Oceania.

### Rare deletion mutants at the *AvrStb6* locus across the species range

We analyzed whether loss-of-function mutants for *AvrStb6* could include gene deletions in addition to premature stop codons. We first inspected the *AvrStb6* region in the species pangenome represented by a set of 19 reference-quality genomes, covering all major wheat producing areas (22). We detected *AvrStb6* homologs in all pangenome isolates except for the Argentinian isolate Arg00. In contrast to the canonical reference IPO323, the Arg00 chromosome 5 appeared truncated near the neighbouring gene downstream of *AvrStb6* (gene_9081 (49); Figure 3), indicative of a complete loss of *AvrStb6*. To assess potential assembly artefacts in the Arg00 genome, we used raw PacBio long reads generated for Arg00 to align against the reference IPO323. The read coverage on chromosome 5 was supporting the fact that both *AvrStb6* locus as well as the neighbouring subtelomeric region were missing (Supplementary Figure S1). Among the 18 pangenome isolates carrying *AvrStb6*, the isolate I93 from Indiana (USA) carried an inverted *AvrStb6* coding sequence without affecting neighbouring genes (Figure 3A). As *AvrStb6* was found missing in a reference-quality genome, we screened for potential additional losses in the thousand-genome panel searching draft genome assemblies. As expected, *AvrStb6* is present in a large majority of isolates, however a small number of isolates showed patterns consistent with deletions such as gene truncation or complete absence of the *AvrStb6* coding sequence. Overall, 34 isolates (3.3%) carried no or no intact *AvrStb6*, of which 23 isolates completely lacked homology and 11 with evidence for a partial deletion (Supplementary Table S6).

**Figure 3:**
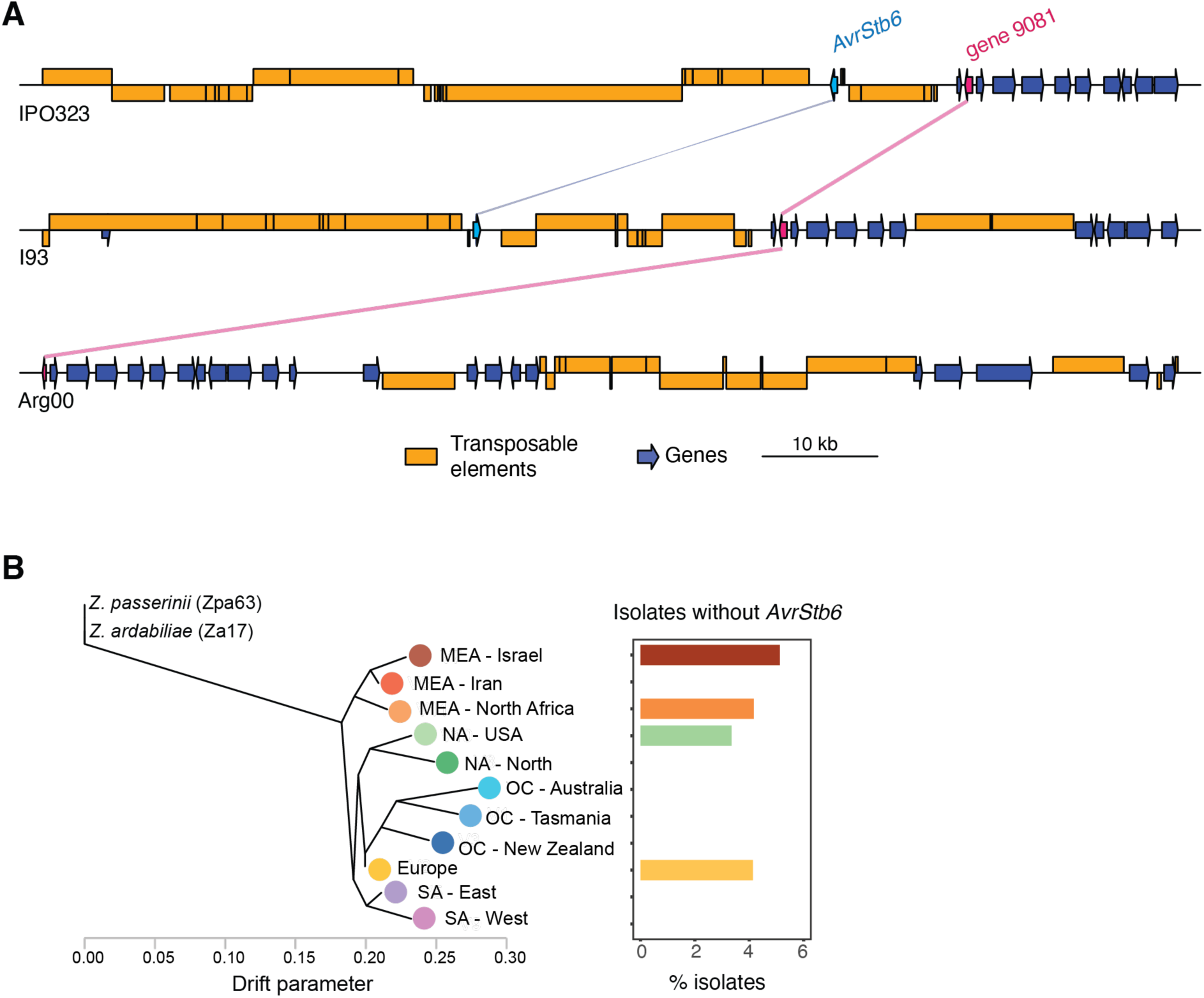
Evidence for *AvrStb6* loss. A) Synteny plot between the telomeric ends and the *AvrStb6* locus on chromosome 5 of the IPO323 reference genome, I93 (USA) and Arg00 (Argentina). Colored lines between chromosomes indicate homologous regions. B) Percentage of isolates missing *AvrStb6* per genetic cluster.

We further investigated evidence for *AvrStb6* loss in the same panel of isolates using copy-number variation (CNV) calls (50). Precision of CNV calling was found to be optimal in 1-kb windows, hence we investigated *AvrStb6* based on the 69-70 kb interval on chromosome 5. Most of the isolates identified as lacking or carrying a truncated *AvrStb6* gene based on draft assemblies (30 out of 34) showed reduced or no sequence coverage in the CNV analyses, as expected (Supplementary Table S6). Isolates without *AvrStb6* were distributed across most genetic clusters (Figure 3B). Next, we analyzed genomes with partial *AvrStb6* sequences to identify potential TE sequences at the synteny breakpoints. We detected two isolates out of 11 with a partial sequence *AvrStb6* sequence and a truncated TE sequence inserted into the locus resulting in a sequence rearrangement. In the European isolate ST16CH_1P7, a DNA/PIF-Harbinger TE fragment was overlapping with the 5’ region of the *AvrStb6* coding sequence replacing a 170 bp segment of the gene. In a second European isolate, 07STF058, a LINE retrotransposon was overlapping with the 3’ end of the truncated *AvrStb6* sequence. Interestingly, this LINE retrotransposon belongs to a family (ZymTri_2023_family_605) detected as one of the physically closest TE families downstream of *AvrStb6* (Figure 2B). Taken together, *AvrStb6* loss and truncation occurs at low frequency in *Z. tritici* populations and at least some of the loss-of-function variants were likely caused by TE-mediated sequence rearrangements.

### Genomic prediction of virulence underpinned by *AvrStb6* variation

Our next objective was to investigate whether *AvrStb6* haplotypes evolved to become more virulent within the species. Gathering phenotypic data from infections is challenging at scale. Hence, we performed genomic predictions parametrized by genome-wide association mapping studies. We used a mapping population consisting of 103 isolates, which was originally designed to identify *AvrStb6* (27). A subset of 87 isolates were overlapping with the thousand-genome panel of the present study. Despite the relatively small GWAS panel size, the genomes encode 10 out of the 59 previously identified *AvrStb6* isoforms, including four of the seven most frequent isoforms. Zhong et al. (27) performed virulence assays on three *Stb6* cultivars Cadenza, Shafir, and Caphorn. Phenotypic readouts included green leaf area percentage, necrotic leaf area percentage, and percentage leaf area containing pycnidia spores in the inoculated area (27). The phenotypic dataset covered a broad spectrum of virulence, ranging from isolates with no lesion induction to complete coverage of leaves by lesions. Taken together, the GWAS panel includes both genetically diverse and globally representative *AvrStb6* haplotypes. Genomic predictions were performed using genomic best linear unbiased predictions (gBLUP). We assessed the consistency of genome predictions by analyzing phenotypes predicted in isolates belonging to the GWAS panel (*i.e.* with associated phenotypic data). Correlation coefficients for observed *vs.* predicted phenotypic values were generally worse using SNPs covering the complete genome compared to focusing the genomic prediction on *AvrStb6*-specific SNPs only (Supplementary Figure S2 and S3). This is consistent with the strong single-locus determinism of *Stb6-AvrStb6* interactions as previously reported (27). Based on the correlation coefficient values, we focused genomic predictions informed by GWAS data on two cultivars and two different phenotypic traits: green leaf area percentage on the Caphorn cultivar and percentage leaf area containing pycnidia spores on the Shafir cultivar.

Genotypes from the center of origin in the Middle East (Iran) were predicted to express the lowest virulence on *Stb6* cultivars, highlighted by the high green leaf area percentage on the Caphorn cultivar (Figure 4A). These isolates showed a higher median green leaf area percentage compared to the avirulent IPO323 isoform (isoform 6) (Figure 4B), suggesting that virulence on these cultivars can be lower than previously reported levels for avirulent isoforms. Pathogen colonization moved from the Middle East to Europe, which represented a steppingstone in the pathogen’s global dissemination to the Americas and Oceania. While most isolates from the European cluster were predicted to express high virulence on *Stb6* cultivars, a wide spectrum of virulence was observed in this cluster, with phenotypic values ranging from 0 to around 80% of green leaf area on the Caphorn cultivar and from 0 to around 60% spore area on the Shafir cultivar (Figure 4A). Isolates from the Oceanian cluster, representing the most recently colonized continent by *Z. tritici*, were predicted to express the highest virulence on *Stb6* cultivars (Figure 4A). Conversely, isolates from the South America East cluster were predicted to be of low virulence on *Stb6* cultivars, especially considering green leaf area percentage on the Caphorn cultivar, expressing similarly low virulence on *Stb6* cultivars as isolates from the center of origin (Figure 4A). Genomic predictions across AvrStb6 isoforms showed that a trend of increasing virulence on *Stb6* cultivars with larger phylogenetic distance to the sister species homolog (Figure 4B). However, genomic predictions for green leaf area percentage on the Caphorn cultivar were showing high virulence in some of the isoforms located closest to the sister species *AvrStb6* homolog (Figure 4B).

**Figure 4:**
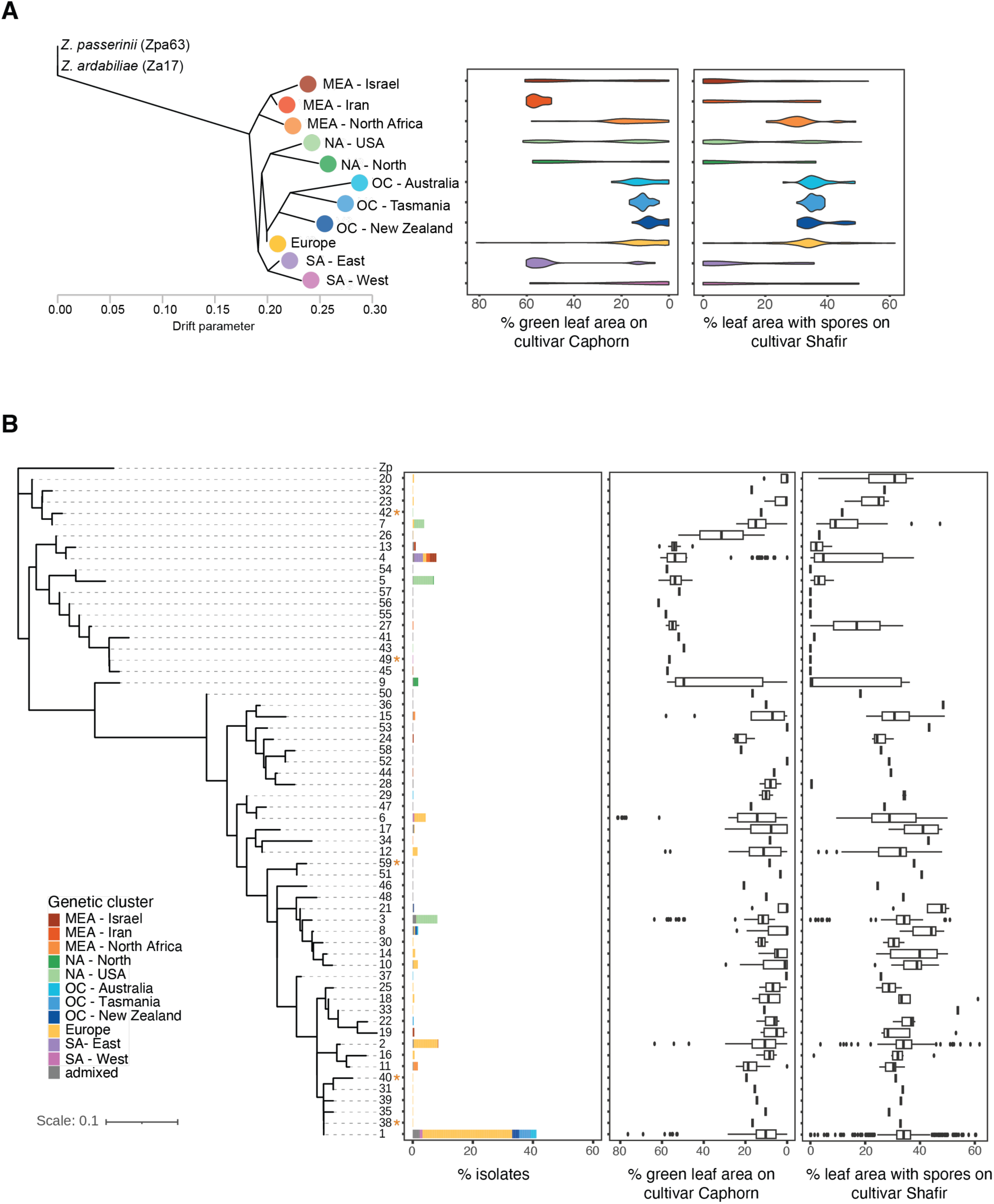
Genomic predictions for virulence on *Stb* cultivars among the *Z. tritici* thousand-genome panel. Predictions were based on including green leaf area percentage in Caphorn cultivar and leaf area percentage covered by pycnidia spores in Shafir cultivar. A) Genomic predictions across population grouped by genetic cluster. B) Phylogenetic tree of AvrStb6 protein isoforms rooted based on the *Z. pseudotritici* (Zp) homolog. Truncated isoforms with premature stop codon are marked with an asterisk. Genomic predictions are summarized by AvrStb6 protein isoform.

Given the global trend of increased virulence on *Stb6* cultivars in more recently established *Z. tritici* populations, we sought to test this pattern at the regional level at high temporal resolution. France is one of Europeans largest wheat producers (59). We genotyped 1327 French wheat cultivars for the presence of *Stb6* using diagnostic markers (56). We then cross-referenced *Stb6* presence with yearly wheat deployment data across the country. The monitoring covered ∼4-5 million hectares representing approximately 80 – 100% of total wheat cultivation in France. *Stb6* prevalence had been widely fluctuating over the 1981-2018 period of the dataset, however there was a marked increase in *Stb6* deployment starting in the mid-1990s up to the most recent years (Figure 5A). Since 2006, *Stb6* cultivars have represented almost half the wheat varieties farmed in France. We investigated whether *Z. tritici* isolates collected in France were shown trends in virulence on *Stb6* cultivars. The available samples covered the period from 1999-2016, a period with steepest increase in *Stb6* resistant allele deployment (from 23% to 51% in 2009, Figure 5A). Genomic predictions for green leaf area percentage on the Caphorn cultivar and leaf area percentage covered by spores on the Shafir cultivar showed only minor changes over years with substantial intra-year variation in predicted virulence trait expression (Figure 5C). Both predicted traits were not significantly explained by the percentage of *Stb6* resistant allele deployment in wheat cultivars across France (regression *R*^2^ ∼ 0; *p*-value = 0.05; Figure 5B). Interestingly, the decrease in virulence observed in 2013 aligns with the fact that most of the French isolates collected that year belong to the MEA–North Africa cluster and not to European cluster as in the other sampled years.

**Figure 5:**
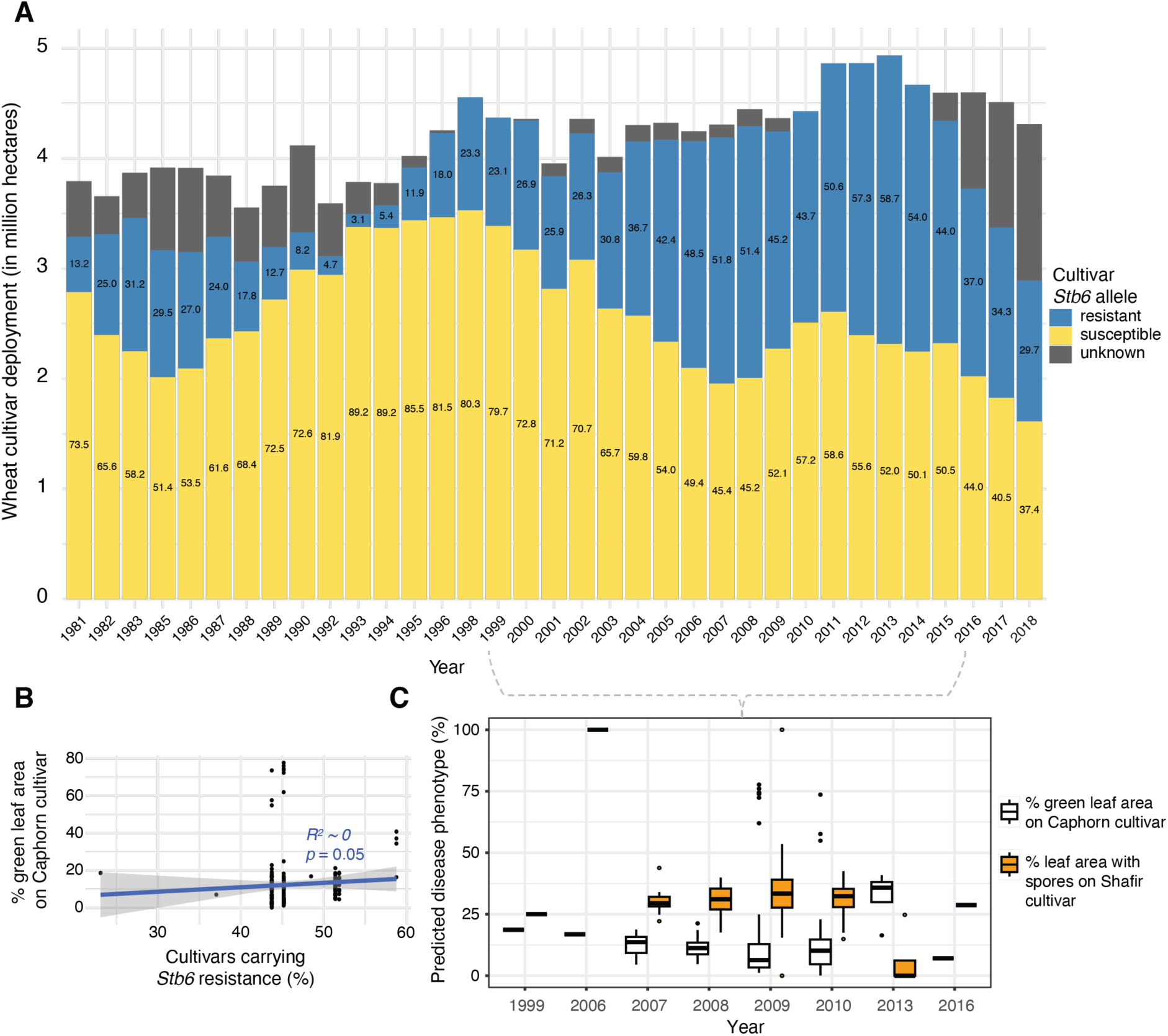
*Stb6* wheat cultivar deployment in France and genomic predictions for *Stb6* virulence. A) Registered wheat cultivar usage (in millions of hectares) as compiled by the DiverCILand database, distinguishing the percentage of cultivars carrying genotyped *Stb6* (resistant or susceptible) alleles. Numbers inside the bars indicate percentages. B) Linear regression of green leaf area genomic prediction as response of the *Stb6* resistance allele deployment in wheat cultivars across France. C) Genomic prediction of *Z. tritici* virulence on *Stb6* cultivars collected in France between 1999 and 2016 including green leaf area percentage on the Caphorn cultivar and leaf area percentage covered by pycnidiospores on the Shafir cultivar.

## Discussion

Plant resistance genes impose high selection pressure on fungal effector genes, causing sequence diversification or even effector deletion. *AvrStb6* is one of the best studied fungal effectors at the population level and interacts in a gene-for-gene manner with *Stb6*, a widely deployed wheat R gene (29). By analyzing a global thousand-genome panel of the pathogen, we show that *AvrStb6* diversified strongly on the European continent with high rates of *Stb6* cultivar deployment. Diversification observation on continents with later *Z. tritici* colonization may have experienced a similar drive by *Stb6* deployment but *Stb6* remains poorly assessed outside of Europe. Consistent with this, European and Oceanian *Z. tritici* populations were also predicted to have evolved higher virulence on *Stb6* cultivars. Sequence rearrangements near effector genes can be important factors driving effector functions. We found that *AvrStb6* is flanked by a dynamic set of TEs with substantial variation across continents and multiple TEs with recent insertion success. TE activity was also the most likely cause for previously unknown *AvrStb6* losses appearing likely independently across the globe.

Rapid effector gene diversification is a convergent pathogen strategy to avoid recognition by the host immune systems (11,60). Previous work on *AvrStb6* (27,32) and other *Z. tritici* effectors including *Avr3D1* (23) reported sequence diversity consistent with strong diversifying selection. Here, we provide a comprehensive view on how effector diversification coincided with pathogen spread at the global scale. *Z. tritici* spread across continents in tight association with wheat cultivation (19,20). In regions of more recent wheat cultivation and more intense production, the deployment of *Stb6*-mediated resistance has been increasing in recent decades (61). Such increased pressure on *Z. tritici* should have caused strong diversifying selection pressure on *AvrStb6* avirulent haplotypes (27,32) including the complete replacement of the originally discovered avirulent haplotypes among most recently collected isolates (33). Within the thousand-genome panel, we have identified 59 AvrStb6 protein isoforms, significantly expanding the previous repertoire of known isoforms (33) with the identification of 37 new isoforms, mainly considering isolates from North America. AvrStb6 isoforms in the Middle East were clustering closest to the homolog in the sister *Z. pseudotritici* homolog, consistent with the center of origin populations having retained among the most ancestral protein variants. European and Oceanian populations carried one of the most frequent but also most divergent isoforms in conjunction with the selection for virulent haplotypes of Avr*Stb6*. The same isoform has also been particularly abundant in recently collected isolates from two continents by Stephens et al. (33).

In *Z. tritici*, as in many other filamentous pathogens, effectors are located in TE-rich compartments, impacting effector and pathogenicity evolution (7,10,62–65). Our investigation of the repetitive region surrounding *AvrStb6* revealed a complex and dynamic landscape of recent TE activity within the species. TE content underwent significant shifts as the pathogen spread from the Middle East to North-Africa and Europe, and subsequently from Europe to the Americas and Oceania. Isolates from the center of origin (Middle East – Israel and Iran) were carrying a MITE DNA transposon and a LINE retrotransposon, as the most frequent TE superfamilies upstream and downstream of *AvrStb6*, respectively. Regions outside the center of origin were predominantly carrying unclassified TEs surrounding *AvrStb6*, except for Eastern cluster in South America. TEs upstream of *AvrStb6* were consistently closer to the effector, whereas those downstream tend to exhibit greater and more variable distance to *AvrStb6* suggesting more frequent sequence rearrangements. Whether the *AvrStb6* locus has more tolerance of upstream insertion activity or whether some insertions might have been favored by selection remains unclear though. The *Z. tritici* effector *Avr3D1* also carries several TEs but of different origin and dynamic (23). The high activity of the MITE DNA transposon was likely a consequence of the weaker apparent defenses and the tendency of MITEs to co-localize with other genes in the *Z. tritici* genome (66). Interestingly, LINEs are completely absent in the *Z. pseudotritici* sister species genome (62), hence their rise to become the most frequent TE downstream of the *AvrStb6* locus in center of origin isolates is striking. *Z. tritici* populations tend to increase TE copy numbers throughout their global colonization history with recently established populations, such as the Americas and Oceania showing marked expansions (20,21). However, in these recent populations, none of the most frequent TE families surrounding *AvrStb6* correspond to the TEs implicated in the strongest TE copy number expansions globally (21,22). This shows that TE insertion dynamics near *AvrStb6* are at least partially independent from genome-wide dynamics and could be governed by selection linked to virulence on *Stb6* cultivars.

The insertion of TEs into coding sequences can result in disruption of transcription or reading frame truncation (67). We identified five isolates (three from Europe and two from the Americas) carrying premature stop codons. However, these mutations were not caused by TE insertions directly but rather by point mutations. Previous evidence of point mutations generating stop codons in *AvrStb6* have been reported by Brunner et al. (32) in an isolate from Oregon (USA). Beyond truncation, we identified the first evidence for low-frequency (∼3%) deletions of *AvrStb6*. In two of the 34 isolates carrying no or no intact *AvrStb6*, gene loss was caused by the insertion of a TE into the coding region. The inserted TEs belonged to different superfamilies, a DNA/PIF-Harbinger transposon and a retrotransposon LINE, showing that the *AvrStb6* loss occurred at least twice independently. Interestingly, the LINE belongs to a family of TEs (ZymTri_2023_family_605) detected as one of the closest TE families downstream of *AvrStb6*, corroborating the idea that genes closer to TEs can be mutagenic (68). *AvrStb6* loss was found distributed across different genetic clusters and continents, reinforcing the idea that the deletion of *AvrStb6* occurred multiple times. In the case of the multihost pathogen *V. dahlia*, TE insertions have also been associated with multiple independent losses of the *Ave1* effector gene, associated with an adaptive response to evade plant host immunity (16). Despite also being located in highly plastic genomic regions, other important *Z. tritici* effector genes, *i.e. Avr3D1* and *AvrStb9*, are not known to be lost (23,69). Earlier work suggested that *AvrStb6* may have an unknown but essential role as no losses were observed (32). However, our findings of multiple, likely independent *AvrStb6* deletions suggest that the gene may be dispensable for survival.

Loss of host resistance following *Stb6* cultivar deployment has been commonly observed in the wheat-*Z. tritici* pathosystem (26,31,70). Using genomic predictions, we were able to construct a global view on virulence expression on *Stb6* cultivars. Our findings suggest that *AvrStb6* virulence has increased as the pathogen evolved in regions where wheat cultivars carrying *Stb6* resistance gene are more widely deployed. This is particularly consistent in Europe with high rates of *Stb6* deployment. European *AvrStb6* diversity has also likely seeded diversity in haplotypes in the Americas and Oceania, allowing for further selection depending on resistance gene deployment levels. For instance, in South America (East), where *Stb6* is not widely used, isolates are predicted to have low virulence on *Stb6* cultivars. On the contrary, in Oceania, isolates are predicted to show the highest levels of virulence on *Stb6*. Using *Stb6* deployment data across France, a major wheat producing country, we found a consistent increase in deployment. The temporal resolution and sampling depth of *AvrStb6* haplotype diversity in France was not sufficient to test for clear associations with cultivar deployment. However, our data is consistent with more virulent *Z. tritici* strains to be favored with increased *Stb6* deployment. Overall, we show that effector locus diversification can occur rapidly and produces complex geographic and temporal patterns within a single plant pathogen species. Rapid sequence evolution of *AvrStb6* spans the spectrum of known effector modifications across different pathosystems including loss of the recognized effector, gains of virulence driven by resistance gene deployment and complex TE insertion dynamics. Our work highlights the power of large genome sequencing panels covering the known distribution range to unravel processes of rapid pathogen adaptation.

## Declarations

### Data availability

Genome assembly data is available from https://doi.org/10.5281/zenodo.13645366. All other data is available from the Supplementary Information or is referenced in the Methods section.

## Acknowledgements

We are thankful to Alice Feurtey for the thousand-genome panel genotyping dataset and Tobias Baril for sharing transposable element annotations. We acknowledge Jonathan Kitt and Sophie Bouchet analyzing wheat cultivars carrying *Stb6* from the 4605 wheat collections. We are grateful to Remi Perronne and Melanie Polart Donat for their contribution to the DiverCILand development. We thank FranceAgriMer for the provision of acreages of bread wheat varieties and Agreste for the total production area of bread wheat.

