## Supplementary Figure for "Diversification, loss, and virulence gains of the major effector AvrStb6 during continental spread of the wheat pathogen *Zymoseptoria tritici*"

Ana Margarida Sampaio *et al*.

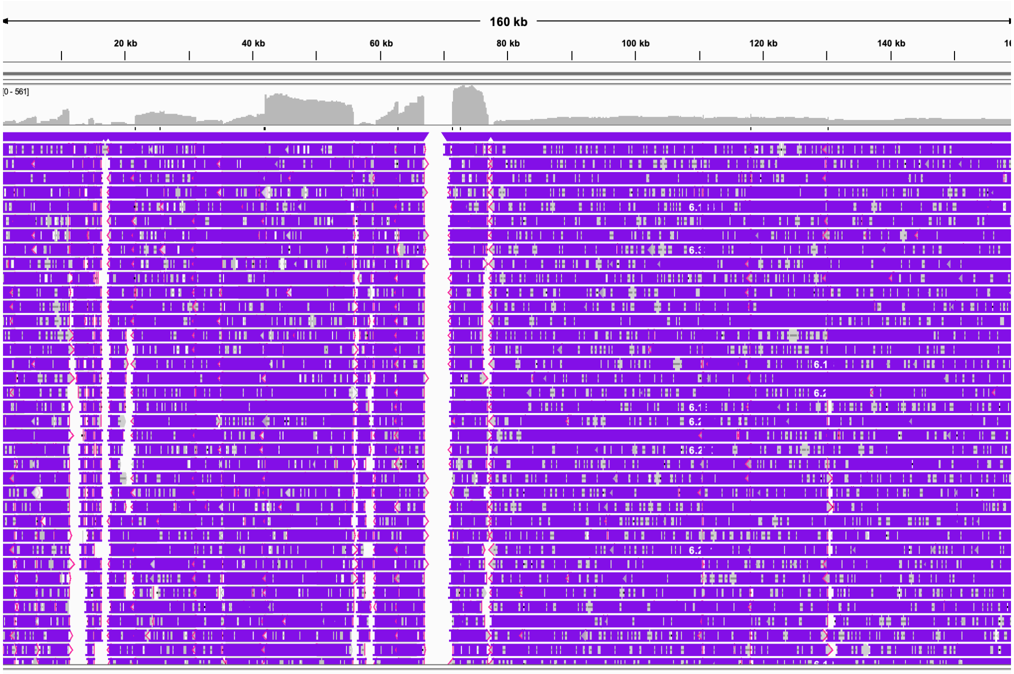


*AvrStb6*

**Supplementary Figure S1**: Visualized mapped PacBio long reads produced from isolate Arg00 mapped to the IPO323 reference genome centered on *AvrStb6*. *AvrStb6* is located at 69019-69383 bp.

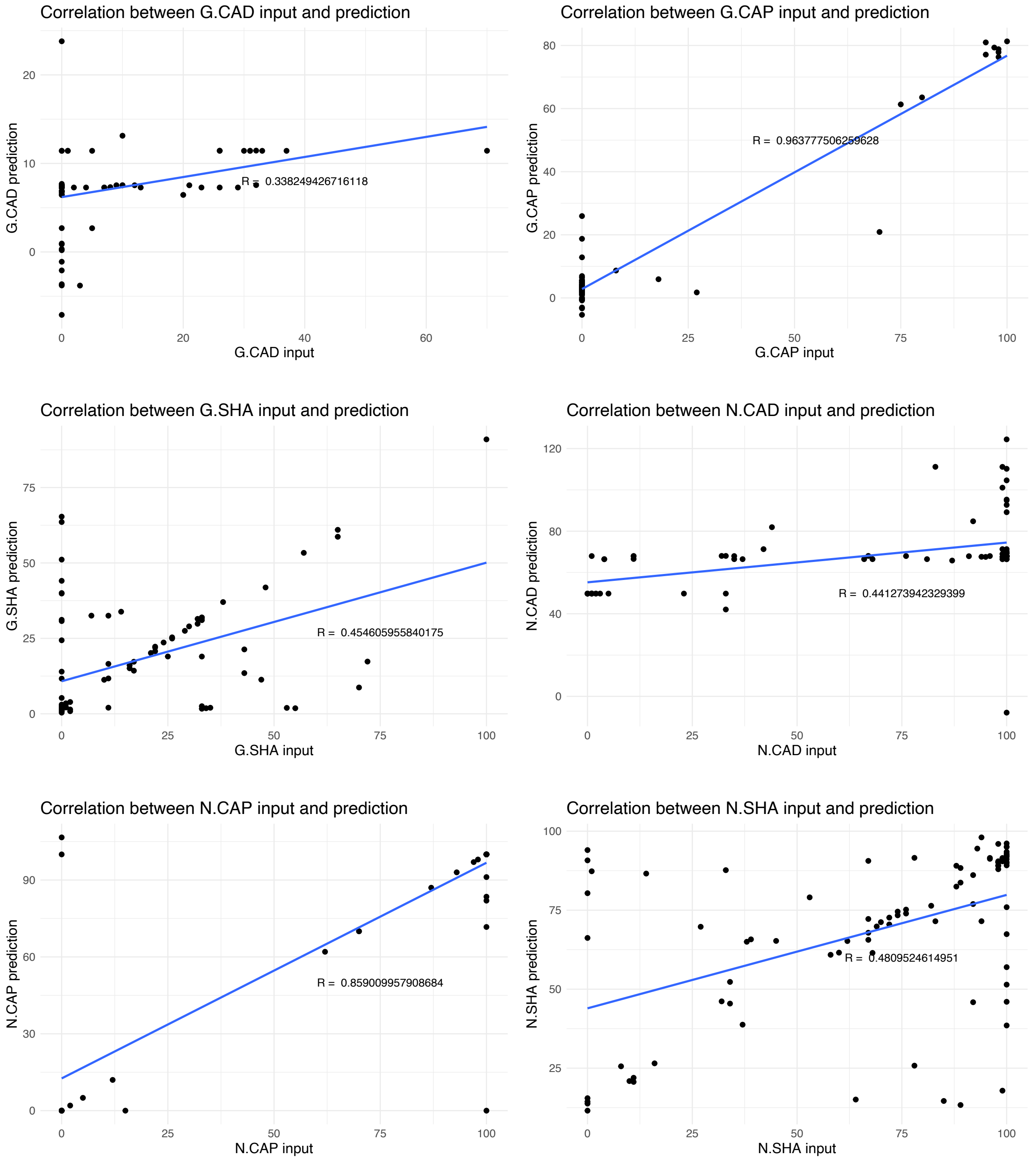


**Supplementary Figure S2**: Correlation plots between phenotypic data collected on the mapping population and predicted phenotypic values based on a SNP matrix covering the *AvrStb6* region and 1000 bp up and downstream of the effector. Correlations were assessed among all three traits (G - green leaf area percentage, N - necrotic leaf area percentage, S - leaf are percentage containing pycnidiospores within the inoculated area) across the three wheat cultivars (CAD – Cadenza; SHA – Shafir; CAP – Caphorn). *R* values indicate the Pearson correlation coefficient between phenotypic data and predicted values for the same isolates.

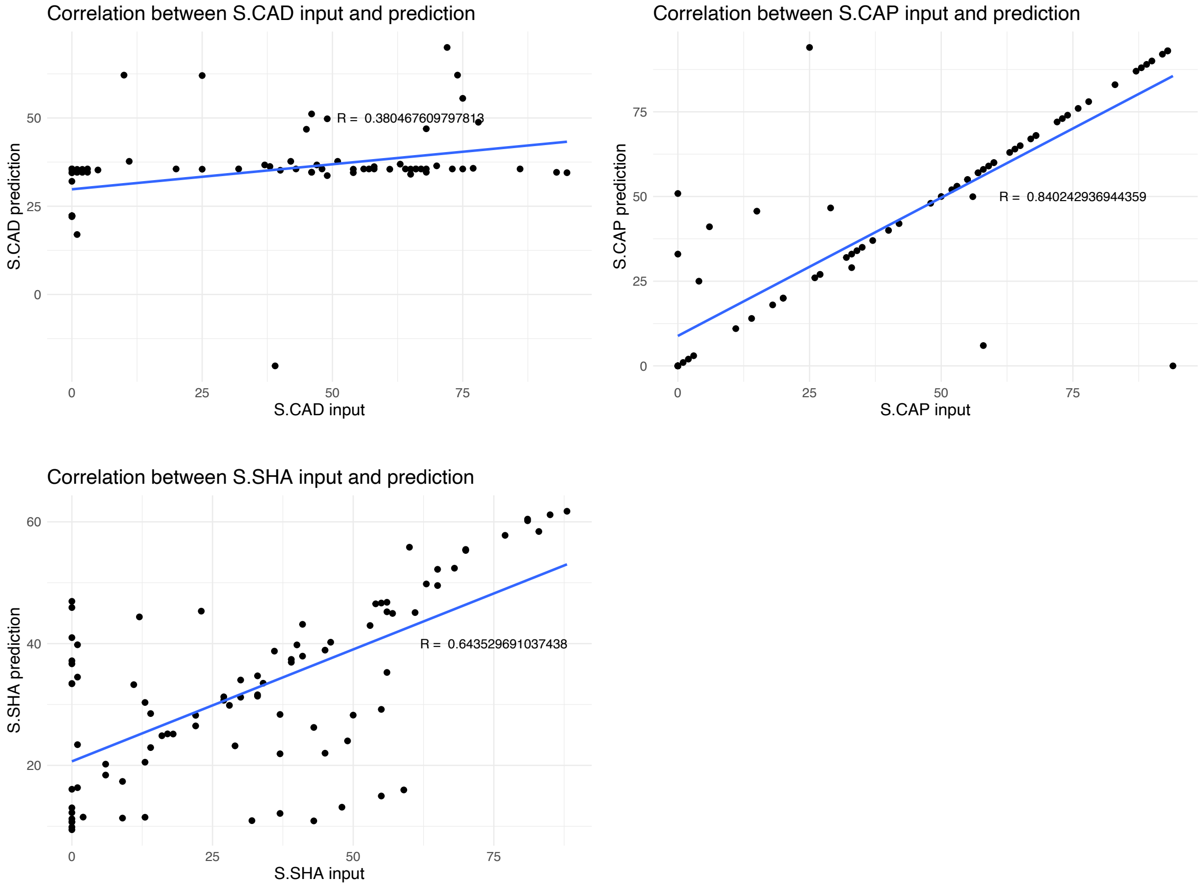


**Supplementary Figure S2** *(continued)*

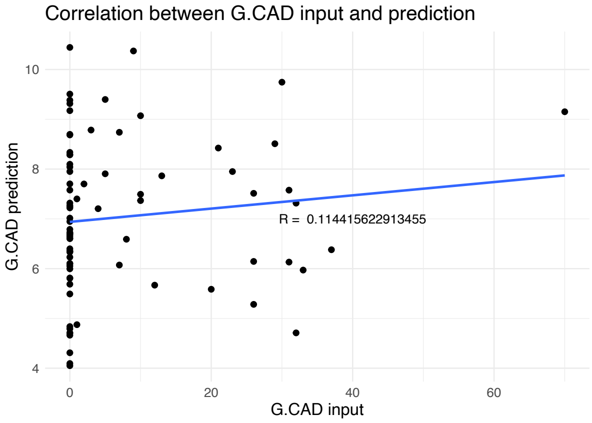
Correlation between G.CAD input and prediction
G.CAD prediction
G.CAD input
R = 0.114415622913455
10
8
6
4
0
20
40
60

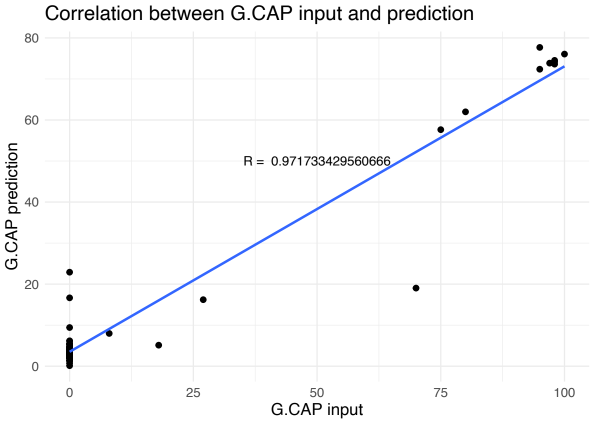
Correlation between G.CAP input and prediction
G.CAP prediction
G.CAP input
R = 0.971733429560666
80
60
40
20
0
0
25
50
75
100

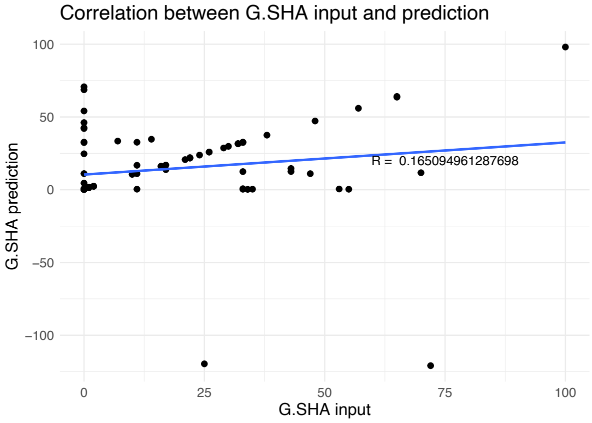
Correlation between G.SHA input and prediction
G.SHA prediction
G.SHA input
R = 0.165094961287698
100
50
0
−50
−100
0
25
50
75
100

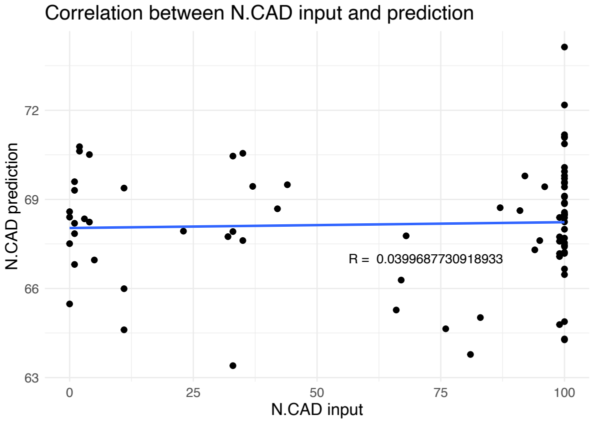
Correlation between N.CAD input and prediction
N.CAD prediction
N.CAD input
R = 0.0399687730918933
72
69
66
63
0
25
50
75
100

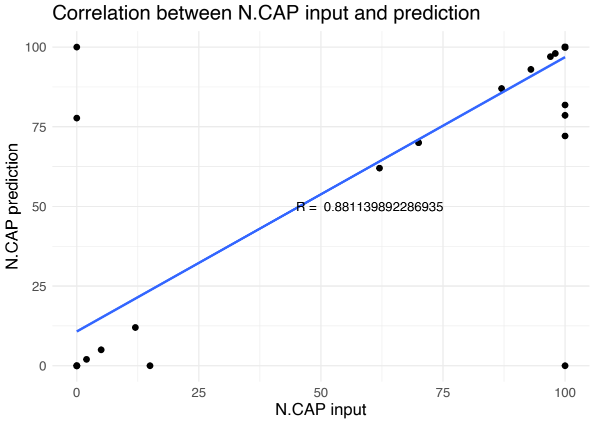
Correlation between N.CAP input and prediction
N.CAP prediction
N.CAP input
R = 0.881139892286935
100
75
50
25
0
0
25
50
75
100

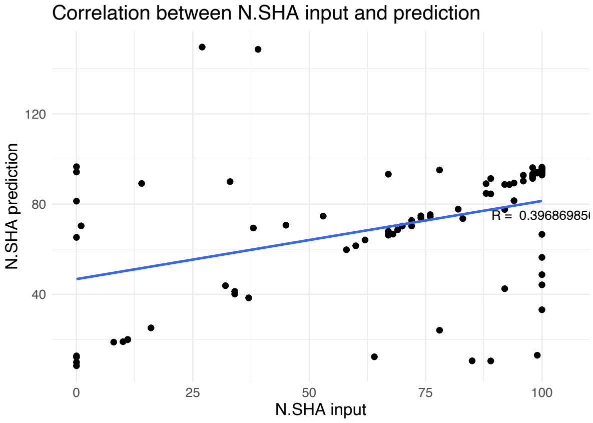
Correlation between N.SHA input and prediction
N.SHA prediction
N.SHA input
120
80
40
0
25
50
75
100

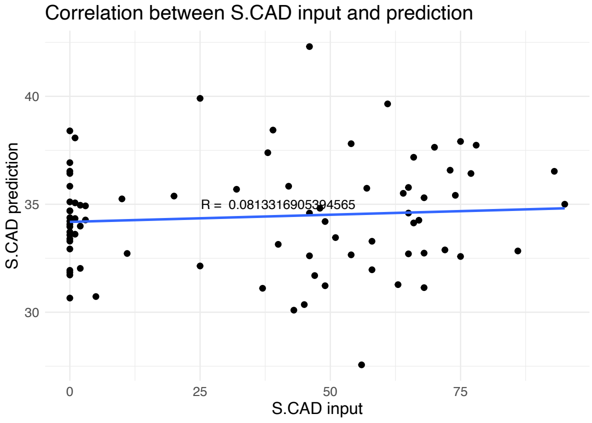
Correlation between S.CAD input and prediction
S.CAD prediction
S.CAD input
40
35
30
0
25
50
75

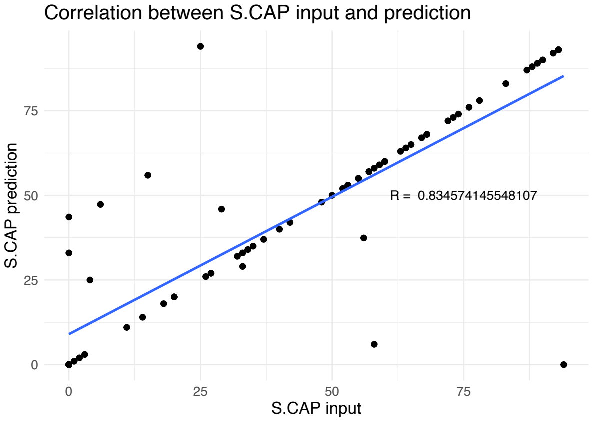
Correlation between S.CAP input and prediction
S.CAP prediction
S.CAP input
R = 0.834574145548107
75
50
25
0
0
25
50
75

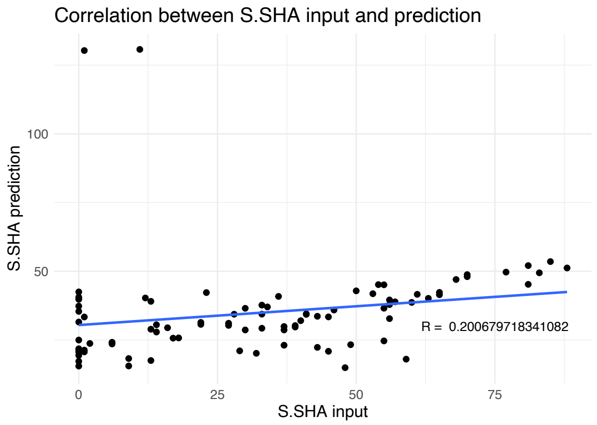


**Supplementary Figure S3**: Correlation plots between phenotypic data collected on the mapping population and predicted phenotypic values based on a SNP matrix covering the entire genome. Correlations were assessed among all three traits (G - green leaf area percentage, N - necrotic leaf area percentage, S - leaf are percentage containing pycnidiospores within the inoculated area) across the three wheat cultivars (CAD – Cadenza; SHA – Shafir; CAP – Caphorn). *R* values indicate the Pearson correlation coefficient between phenotypic data and predicted values for the same isolates.
